# *Ex vivo* dynamics of human glioblastoma cells in a microvasculature-on-a-chip system correlates with tumor heterogeneity and subtypes

**DOI:** 10.1101/400739

**Authors:** Yang Xiao, Dongjoo Kim, Burak Dura, Kerou Zhang, Runchen Yan, Huamin Li, Edward Han, Joshua Ip, Pan Zou, Jun Liu, Ann Tai Chen, Alexander O. Vortmeyer, Jiangbing Zhou, Rong Fan

## Abstract

The perivascular niche (PVN) plays an essential role in brain tumor stem-like cell (BTSC) fate control, tumor invasion, and therapeutic resistance. Herein we report on the use of a microvasculature-on-a-chip system as a PVN model to evaluate the dynamics of BTSCs *ex vivo* from 10 glioblastoma patients. We observed that BTSCs preferentially localize in the perivascular zone. Live cell tracking revealed that the cells residing in the vicinity of microvessels had the lowest motility, while a fraction of cells on the microvessels unexpectedly possessed the highest motility and migrated over the longest distance. These results indicate that the perivascular zone is a niche for BTSCs, while the microvascular tracks are also a path for long-distance tumor cell migration and invasion. Additionally, the degree of co-localization between tumor cells and microvessels varied significantly across patients. To validate the results from our microvasculature-on-a-chip system, we used single-cell transcriptome sequencing (10 patients and 21,750 single cells in total) to identify the subtype of each tumor cell. The co-localization coefficient was found to correlate positively with proneural (stem-like) or mesenchymal (invasive) but not classical (proliferative) tumor cells. Furthermore, we found that a gene signature profile including PDGFRA correlated strongly with the “homing” of brain tumor cells to the PVN. Our findings demonstrated that *ex vivo* dynamics of human brain tumor cells in a microvasculature-on-a-chip model can recapitulate *in vivo* tumor cell dynamics, heterogeneity, and subtypes, representing a new route to the study of human tumor cell biology and uncover patient-specific tumor cell functions.

## INTRODUCTION

The brain tumor perivascular niche (PVN)(1, 2), the region in the vicinity of microvessels is a prime location for brain tumor stem-like cells (BTSCs). Tumor microvasculature creates a complex microenvironment(3) consisting of various cell types, the extracellular matrix, and soluble factors that mediate cell-cell interaction. The brain tumor PVN controls maintenance, expansion, and differentiation of BTSCs via direct cell contact or paracrine signaling cues(4-6). BTSCs often receive bidirectional crosstalk from endothelial cells and other cell types in the niche(7). In addition, the perivascular zone is a path for tumor cells to migrate over long distances. Unlike other solid tumors, glioblastoma multiforme (GBM) cells rarely metastasize to other organs, but they can invade the entire brain by migrating along specific brain tissue structures, such as blood vessels or white matter tracts, leading to high rates of relapse(8-11). Despite the success in modeling diffuse brain tumors in both genetically-modified and patient-derived xenograft (PDX) animals(12), there is an unmet need for an *in vitro* system that can bridge conventional cell culture and animal models by mimicking not only the anatomy but also the function of the PVN to study the dynamics of BTSCs.

Traditional 2D cell cultures are incapable of replicating *in vivo* 3D environments where cancer cells reside, and may result in inaccurate data to evaluate drug responses(13, 14). Thus, there have been substantial efforts to develop 3D cell culture models, as well as patient-derived tumoroids, that exhibit features closer to *in vivo* conditions(15, 16). These include areas of hypoxia, heterogeneous environment (e.g., stromal cells), different cell proliferation zones (quiescent vs. replicating), ECM influences, soluble signal gradients, and differential nutrient and metabolic waste transport(17-20). Existing techniques used to culture cells into 3D structures include scaffold-based approaches (e.g., polymeric hard scaffolds, biologic scaffolds, micropatterned surfaces)(21-23) and non-scaffold-based approaches (e.g., hanging drop microplates, spheroid microplates containing Ultra-Low Attachment (ULA) coating, and microfluidic 3D culture)(20, 24, 25). To date, current *in vitro* cancer models lack perfusable microvasculature, and thus, may not capture the essential role of microvascular niches in tumor progression and therapeutic response(26-28).

Recently, perfusable microvasculature in microfluidic systems have been developed(29-38). Several studies reported a spontaneous microvasculature formation via a vasculogenesis-like process in a hydrogel loaded microfluidic chamber(29, 30), in which the seeding of endothelial cells in hydrogel-loaded microfluidic networks leads to endothelial proliferation and lumen formation(31-34). Kamm et al. developed a microfluidic device containing 3D perfusable microvessel networks to investigate tumor cell intra- and extravasation(35, 36). Phan et al. reported 3D microtumors/tumoroids made of colorectal, breast, and melanoma tumor cell lines can be re-vascularized in a perfusable diamond-shaped microchambers to study drug responses(39). To our knowledge, current tumor-on-a-chip models incorporated cancer cell lines as a proof-of-concept, and it has not yet been demonstrated that such a platform can work as a pathophysiologically relevant system to evaluate the function of primary tumor cells *ex vivo* in a patient-specific manner.

We report on the first demonstration that a tissue-engineered microvasculature-on-a-chip system can be utilized to examine the function of primary patient-derived BTSCs. Live cell imaging of tumor cell dynamics and localization allowed for delineating the interaction of single brain tumor cells with the nearby microvessels. We found that BTSCs preferentially localize in the PVN, possibly due to their stemness or invasiveness. We further validated our results using single-cell RNA sequencing of 10 glioblastoma patients (26 batches, 21,750 single cells). We found that tumor-microvessel co-localization was based on the genetic and pathologic subtypes of the tumor samples. Our GBM-microvasculature-on-a-chip model demonstrates the potential for *ex vivo* analysis of tumor cell functions and patient-specific glioblastoma treatment.

## RESULTS AND DISCUSSION

### Perfusable endothelialized microvasculature-on-a-chip

Our microfluidic device (AIM Biotech) is comprised of a center chamber for loading a mixture of endothelial cells and hydrogel precursor and two lateral channels for perfusing culture medium (Fig. 1a). The cell/hydrogel loading chamber and two lateral channels are separated by triangular microposts(40) which prevent gel leaking. Human umbilical vein endothelial cells engineered for expression of green fluorescence protein (GFP-HUVECs, Angio-Proteomie) were suspended in a 2.5 mg/ml fibrin gel precursor and loaded into the microfluidic chamber. Premature nascent lumens developed in 3 days via vasculogenesis and a connected microvessel network spanning the entire chamber (∼1.3 mm × 8mm) were established in 4-6 days (**Supplementary Fig. 1a&b**) upon daily perfusion with medium containing a cocktail of growth factors (VEGF(50ng/ml), FGF(20ng/ml), EGF(20ng/ml)). Anastomosis of microvessels was observed at the vessel openings to medium flow channels, which allowed for leak-free continuous perfusion and the establishment of shear stress in the microvascular network to foster microvessel development and maturation. Adding cells that secrete proangiogenic factors in the hydrogel precursor accelerated vasculogenesis and expedite the formation of interconnected perusable microvascular network in as short as 3 days(41).

**Figure 1.**
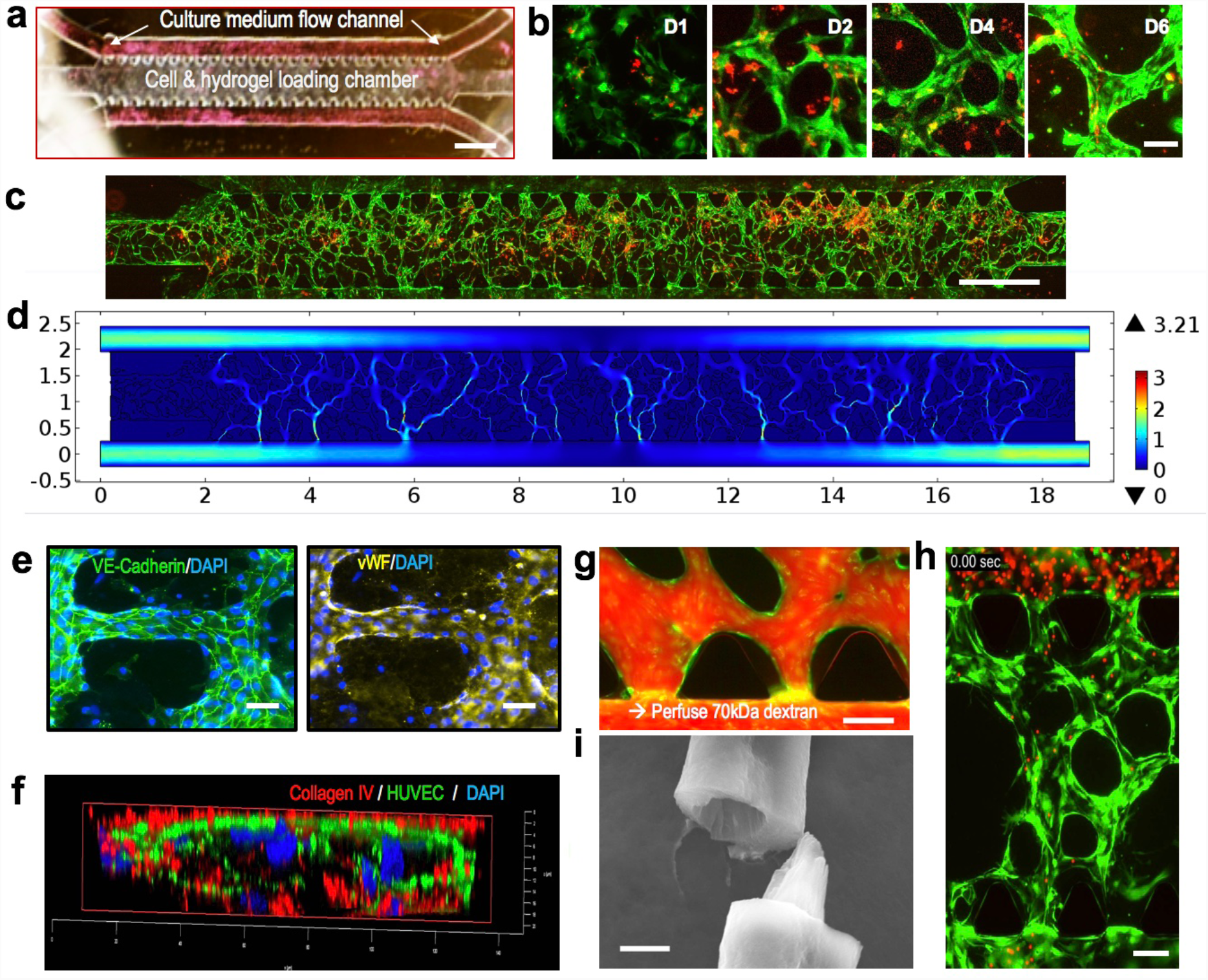
Growth of BTSC-incorporated microvasculature-on-a-chip. (**a**) Microfluidic device (250 μm in height) containing a cell/gel loading microchamber (8000 μm ×1300 μm) flanked by two medium flow channels (500 μm in width). An array of triangular microposts separate the gel chamber and the medium flow channel, allowing for loading and confining the hydrogel precursor to the mid-chamber only. Scale bar: 1000 μm. (**b**) Representative time course images of the microvessel formation over a period of 6 days. Scale bar: 15 μm. Green: GFP-HUVEC. (**c**) Whole chip scan showing microvasculature formation (96 hours post cell in fibrin) and loading of single BTSCs (GS5). Green: GFP-HUVECs. Red: BTSCs. Scale bar: 1000 μm. (**d**) Comsol Finite Element Simulation of flow velocity magnitude (mm/s). The finite element model was constructed using the experimental whole-chip microvessel network in (c). (**e**) Immunostaining of VE-cadherin or vWF to examine the formation of tight junction and the function of tissue-engineered endothelial vessels. Scale Bar: 20 μm. (**f**) Cross-sectional confocal image showing two adjacent microvessels and collagen IV deposition. Green: GFP-HUVECs. Red: collagen IV. (**g**) Infusion of 70-kDa fluorescent dextran to measure impermeability of the lumen and examine microvessel opening to the media channel (anastomosis). Green: GFP-HUVECs. Red: Dextran. Scale Bar: 100 μm. (**h**) Flowing fluorescent microbeads (red) (10um) through a microvessel network (green) that were grown for 3 days. Scale Bar: 100 μm. (**i**) SEM image of the microvessels. Scale Bar: 5 μm.

### Incorporating primary glioblastoma cells in the microvasculature-on-a-chip system

To investigate the behavior of BTSCs in the microvascular environment, we used a well-characterized, patient-derived brain tumor neurosphere culture, GS5 (**Supplementary Figure 2 a&b**), which is enriched for tumor stem-like cells and can generate highly penetrating tumors intracranially in mice(27, 42). GS5 cells were mixed with GFP-HUVECs in the hydrogel precursor at a 1:4 ratio and loaded into the microfluidic device to grow 3D microvasculature. Endothelial cells began to connect together to form networks by day 2 and grew into extensive microvessels by day 4 (Figure 1b). Tumor cells exhibited motility while the microvessels were remodeling until reaching a relatively stable geometry by day 6.

**Figure 2.**
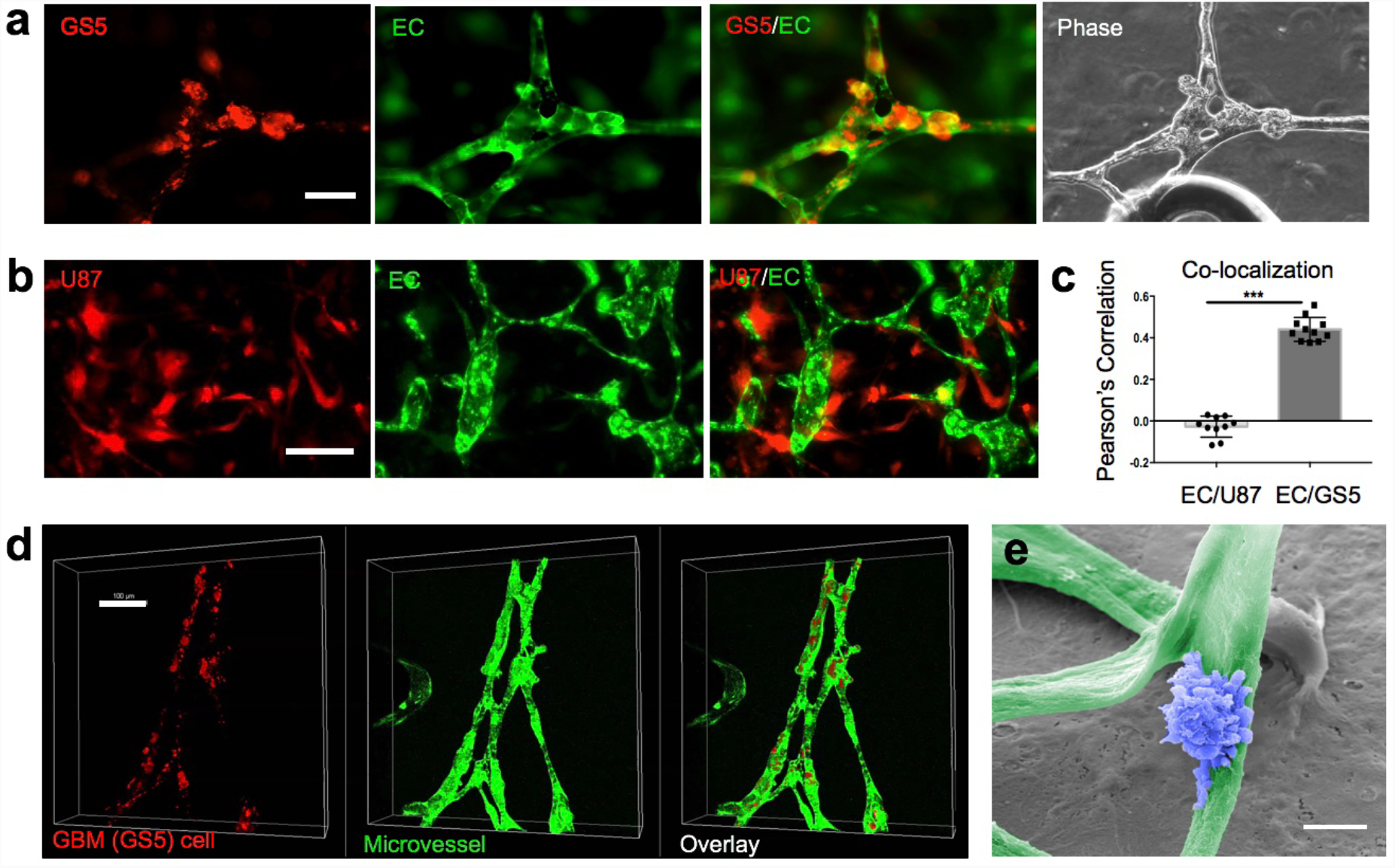
Quantification of tumor cell localization relative to microvessels. (**a**) Phase and fluorescent images of microvessels with BTSC GS5. BTSCs were incubated with the Dil membrane dye for 40min prior to co-culture with GFP-HUVECs. BSTCs localize more preferentially to the branching points of the microvessel network. Scale bar: 80 μm. (**b**) Fluorescent images of glioblastoma cell line RFP-U87 cells loaded in a microvessel network. RFP-U87 cells randomly distributed in the gel space and the microvessels constantly remodeled. Scale bar: 80 μm. (**c**) Quantitative analysis of co-localization of GS5 vs U87 cells with the microvessels. (**d**) Confocal images to examine co-localization of GS5 cells and microvessels. Red: GS5 cells. Green: HUVECs. Scale Bar: 100 μm. (**e**) SEM image (false color) showing a BTSC on the microvessel. Scale:5 μm.

A whole chip scan (Figure 1c) showed the formation of interconnected microvessel network in day 4. After determining the patency of the microvessel network, finite element simulations were performed to determine the local flow rates (Figure 1d) and shear stresses in the microvessel bed. Florescent images were vectorized and imported into COMSOL Multiphysics to simulate the flow of media. With a gradient of 10Pa, the maximum shear stress experienced by microvessels was 4.02 dyne/cm^2^ and the maximum flow rate in the microvessels was 3.21mm/s. The simulations were able to reveal the heterogeneity that exists within the microvessels to further evaluate whether flow and shear properties influence the interaction between tumor cells and the adjacent vessels.

To confirm that tissue-engineered microvessels are physiologically functional and resemble native microvessels, we examined the formation of lumen and the presence of protein markers indicative of essential physiological characteristics. Confocal imaging showed that our microvessel network developed open hollow lumens as early as day 3 (**Supplementary Figure 1c**). Immunostaining of the fixed microvessels for VE-Cadherin revealed the formation of endothelial cell tight junctions over a large area of the microvascular network (Figure 1d **left**). In response to shear stress induced by medium perfusion, microvessels produced von Willebrand factor (vWF), a secreted factor that was deposited onto the inner wall of endothelia and polymerized to form vWF fibers that can facilitate platelet adhesion (Figure 1d **right**). Cross-sectional confocal imaging also revealed that endothelial cells secreted and deposited collagen IV to the basal surface of lumen, forming a 3D collagen IV mesh as early as day 3. This collagen layer not only stabilizes microvessels but also constitutes an important ECM component of the PVN in our model (Figure 1f). This also demonstrated endothelial cell polarization as they formed lumens. In addition, the production of collagen IV and deposition of ECM are indicative of microvessel maturation.

To further evaluate the quality of microvessels, we tested the permeability via the perfusion of 70kDa fluorescently labeled dextran (Figure 1g). We used time-lapse imaging to calculate a permeability coefficient of (6.76±0.92) × 10^−7^ cm s^−1^ (n=3, at day 6), which was within the range of previously reported values for *in vitro* microvessel models(30, 41). Furthermore, a suspension of 10μm-sized fluorescent polystyrene beads was perfused into the upper channel of a 3-day-old chip. We observed that the beads readily traveled through the microvascular network and entered the lower microchannel with minimal adherence to the microvessel wall (Figure 1h and **Supplementary Movie 1**). Finally, the microvasculature hydrogel slab was retrieved, fixed, and fractured to expose the cross-section of microvessels, which was imaged with scanning electron microscopy (SEM) to confirm the formation of hollow lumen (Figure 1i).

### Preferential localization of BTSCs in PVN

The role of PVN in controlling BTSC fate has been reported in human glioblastomas and validated with animal xenograft models(4, 5, 43, 44). Using tissue-engineered microvasculature models to determine whether BTSCs preferentially localize within the PVN, we quantified co-localization of microvessels and BTSCs (GS5) relative to a glioblastoma cell line (U87). Tumor cells were pre-stained with lipophilic cell tracking dye Dil (Invitrogen), mixed with GFP-HUVECs, and loaded into the microfluidic chip to examine microvessel growth and tumor cell dynamics. After 7 days, we observed that BTSCs preferentially localized in the perivascular zone (Figure 2a), specifically in the bifurcation region of the microvessel network. In contrast, U87 cells did not co-localize in the perivascular zone (Figure 2b). In addition, we observed that perfusion of U87 cells led to constant microvessel remodeling and unstable microvessel network, whereas GS5 cells resulted in stabilized microvessel network in 4-5 days. Quantitative analysis confirmed that patient derived BTSCs (GS5) showed a significantly higher Pearson’s correlation coefficient (0.44± 0.02, n=11) than U87 cells (−0.03 ± 0.02, n=10) (Figure 2c) 7 days after loading into the microchip.

Scanning confocal microscopy was used to show the 3D location of BTSCs (red) relative to microvessels (green) (Figure 2d). The distribution of BTSCs in this 3D hydrogel slab mirrored the structure of the microvessel network. We did not observe tumor/endothelial cell co-localization where the perfusable lumens did not form with U87. Furthermore, the BTSCs appear to be fully integrated in the endothelia of microvessels. It has been reported that brain tumor stem cells can differentiate into vascular cells, such as pericytes or endothelial cells, and contribute to tumor angiogenesis(45-47). We found that most tumor cells adhere and spread onto the surface of microvessels, presumably via adhesion to collagen IV mesh produced by endothelial cells (Figure 1e). In contrast to the smooth surface of endothelialized microvessels (**Supplementary Figure 2c**), BTSCs appear to be very rough in cell surface and are decorated with extensive vesicles (**Supplementary Figure 2d**), suggesting their secretory activity to elicit a more complex cell-cell communication network in the PVN(48, 49).

Our findings indicate the potential of a tissue-engineered microvasculature-on-chip system as a functional surrogate to examine the interaction between tumor cells and the PVN, and potentially as a platform for *ex vivo* assay of tumor cell properties. Although the underlying mechanisms remain unclear, we believe they are related to the signals associated with oxygen gradients, nutrient gradients, endothelial cell-secreted factors, and interactions between BTSCs and ECM on the vessel surface.

### Tracking tumor cell migration in PVN

We observed extensive co-localization of GS5 BTSCs in the perivascular region 7 days after the formation of interconnected microvessel network (Figure 3a). However, how each cancer cell migrates relatively to microvessels to eventually “home” in the perivascular region remains unclear. To explore the underlying mechanisms, we utilized live cell tracking fluorescence microscopy (Figure 3b) to image GS5 cells in the microvasculature-on-a-chip device 2 days after cell seeding for a period of 20 hours (48-68Hr post cell-loading) at a rate of two scans per hour. We observed that most tumor cells (i.e., cell 3 in Figure 3b) residing in the perivascular region had reduced migration rates and exhibited a round shape morphology. The cells more distant from the microvessels showed spindle-like morphology and actively extended filopodia to sense the surroundings but did not migrate over long distances. Surprisingly, the most migratory cells (i.e., cell 1) were located very close to or traveled along the microvessels. It is known that brain tumor cells utilize the microvascular tracks to invade distant regions of the brain. The migratory phenotype is predominately de-differentiated or mesenchymal, resembling the GBM mesenchymal subtype that often causes non-resectable disease(50-52). Our data is consistent with prior *in vivo* studies(11) and suggests that our device is capable of differentiating between highly invasive tumor cells and stem-like quiescent tumor cells.

**Figure 3.**
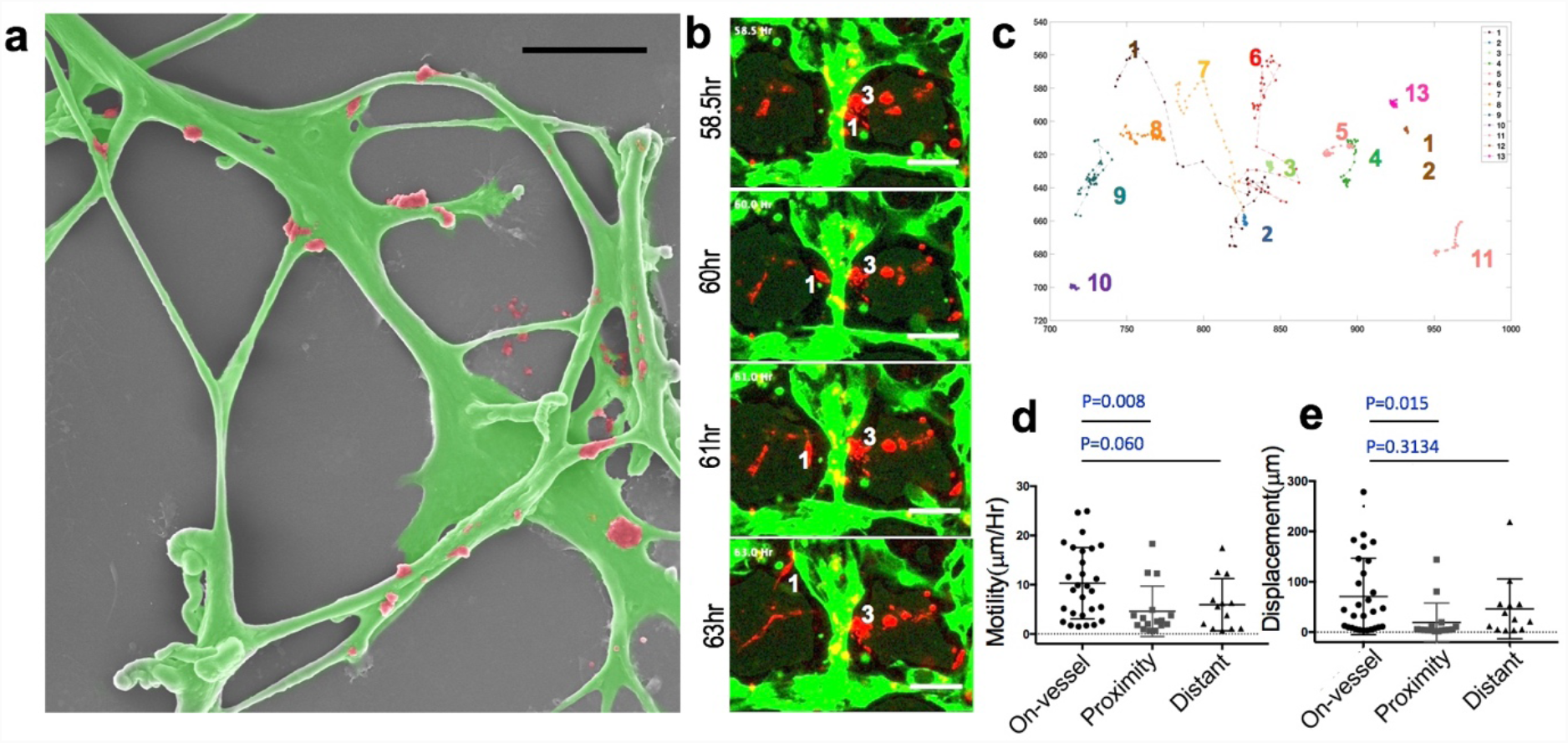
Tumor cell migration in the microvasculature-on-a-chip. **(a**) SEM image (false color) of a typical microvessel network (green) with GS5 cells (red). Scale:50μm (**b**) Fluorescence images of a representative region at different time points to track tumor cell migration. (**c**) Migration trajectory of 13 tumor cells in the region shown in (b) for a day 3 microchip and measured over a period of 20Hr. (**d**) Average motility (μm/Hr) of single tumor cells from three groups defined by the initial location of tumor cells relative to microvessels. On-Vessel (10.29±7.23 μm, n=28); Proximity (4.60 ± 5.11 μm/Hr, n=16); Distant (5.94± 5.29 μm/Hr, n=13). ANOVA test (p=0.0133, Kruskal-Wallis Test). (**e**) Absolute displacement of single tumor cells from three groups. On-Vessel (70.71±75.96 μm, n=28); Proximity (19.32 ±38.42 μm, n=16); Distant (46.28± 59.5 μm, n=13). ANOVA test (p=0.0042, Kruskal-Wallis Test).

Next, we calculated cell motility and migration distance to determine the cell migration trajectory (Figure 3c, **Supplementary Figure 3, and Supplementary Movie 3**). We grouped the cells based upon each cell’s relative distance to the nearest microvessel into three categories: *On-Vessel* (the distance to vessel in our fluorescence image = 0), *Proximity* (the distance to vessel < 50um) and *Distant* (the distance to vessel >= 50um which is considered as outside the PVN). Statistical analysis (One Way ANOVA) demonstrated significant differences in the total displacement (p=0.0121) or motility (p=0.0134) (Figures 3d&e) among these groups. Tumor cells residing in the PVN were mostly round but the ones outside the PVN were constantly extending filopodia to explore their surroundings. Interestingly, the lowest migratory distance cell group was not “On Vessel” but the “Proximity” group. The “On Vessel” group was up to 5x higher motility compared to “Proximity” group. We hypothesized that the direct interaction between tumor cells and the collagen mesh on the vessel surface was responsible for facilitating tumor cell adhesion and directional migration.

The surface marker phenotype was also measured by immunostaining for nestin and So×2 and the result confirmed that a significant fraction of brain tumor stem-like cells was perivascular. In the PVN, 61.6% ± 2.2%(n=3, day 7) of BTSCs were nestin-positive cells and 68.2 ± 4.0% positive for Sox2. The observation of tumor cell differential dynamics in PVN was unanticipated and demonstrated the feasibility to use live cell dynamics measured in our device to assay the functional phenotype of single brain tumor cells in a physiologically relevant environment.

### All patient samples: co-localization of tumor cells and microvessels

To assess inter-patient heterogeneity of brain tumor cells’ PVN “homing” ability, we applied the same approaches to evaluate additional 9 patients’ BTSCs. All patient samples (Figure 4a) used were IDH wild type, glioblastomas, with differing MGMT promoter methylation, EGFR amplification status, stem-cell markers So×2 or nestin, (Figure 4b), and ability to grow tumors in mice. A representative whole chip scan (sample GBM6, Figure 4c) showed extensive microvessel growth by day 4, while BTSCs largely remain isolated. The relative distance between each tumor cell and the nearest microvessel was measured to calculate a Pearson’s co-localization coefficient R value for each patient sample. The results together with GS5 cell data in Day 4 were rank ordered and plotted in Figure 4d. GS5 ranked highest, and other top ranked patient samples included GBM6, GBM24 and GBM5. The lowest co-localization coefficient in the group was patient GBM12, with a co-localization coefficient nearly a third of that for GBM6. We noticed significant morphological differences in tumor cells when interacting with microvessels (Figure 4e). For example, the typical perivascular GBM5 cells wrapped around the microvessels and actively produced microvesicles. The morphology of GBM6 cells in the microvessel network resembled that of brain tumor microvasculature in murine(11).

**Figure 4.**
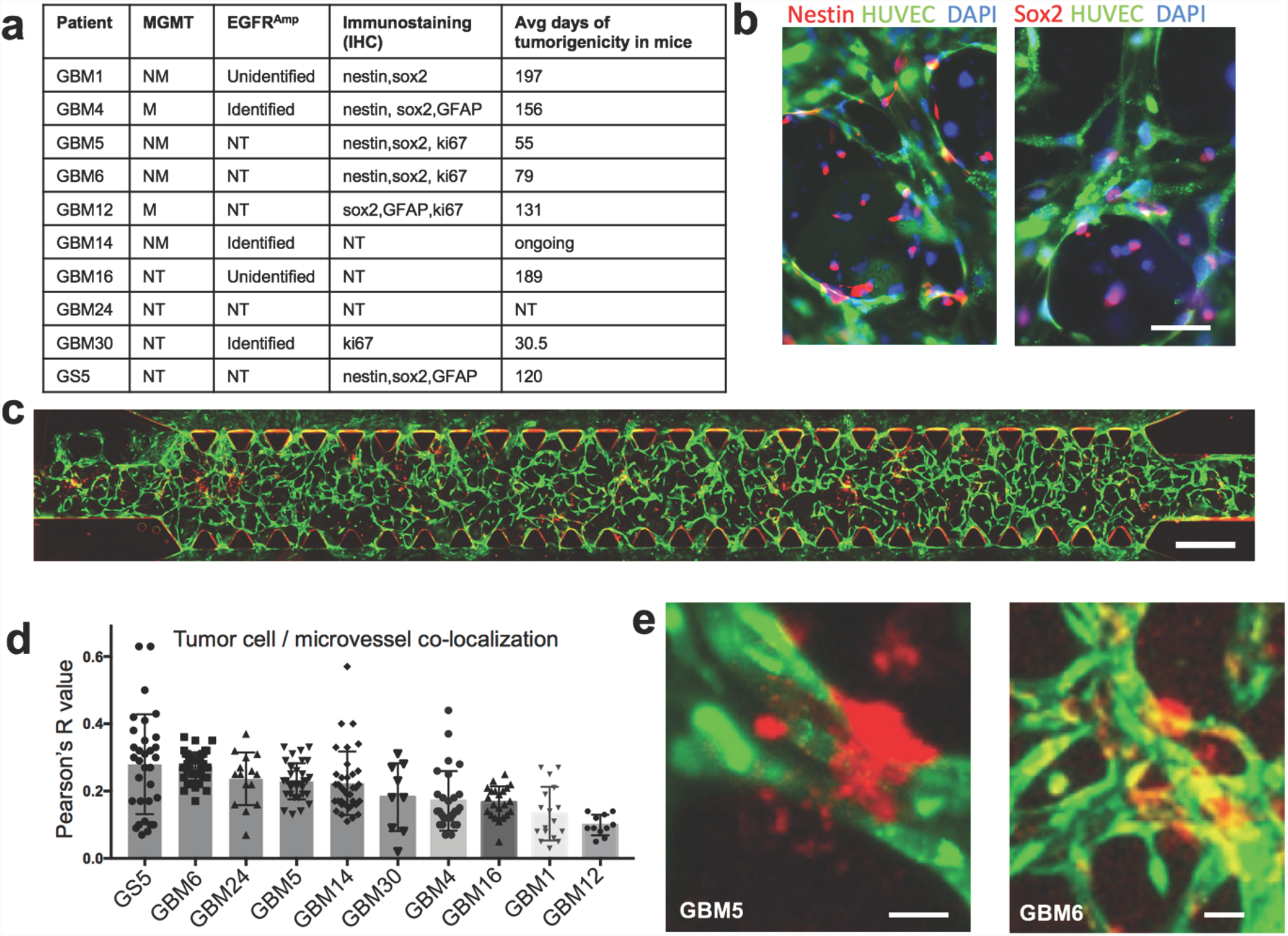
All patient samples: tumor cell localization relative to microvessels. (**a**) Patient information table. All GBM samples are IDH^wt^. M=promoter methylated, NM=promoter not methylated, NT=not tested. (**b**) Immunostaining of nestin and Sox2. Scale: 30 μm. (**c**) Whole chip scan of GBM6 in the microvasculature chip at Day 4. Scale: 700 μm. Red: GBM6 cells. Green: HUVECs. (**d**) Co-localization coefficient of tumor cells and microvessels measured for all patient samples. (**e**) Representative images of patient cells in the microvasculature chip. Red: GBM cells. Green: HUVECs. Scale: 10 μm.

### Single-cell RNA-seq to correlate with transcriptional subtypes and gene signatures

To correlate the observed brain tumor cell behavior heterogeneity with the microvasculature-on-a-chip system to tumor cell genotype or phenotypes, we conducted single-cell 3’ mRNA sequencing of all ten patient samples (**Supplementary Table 1**) and obtained 26,027 single-cell transcriptomes at a depth of at least 10,000 reads per cell. To minimize the sequencing batch-to-batch bias, we prepared 2-3 batches of cDNA libraries for each patient sample for a total of 26 batches. The mean of transcripts (UMIs) per cell detected in each batch ranged from 6192 to 20174. The median of genes detected per cell ranged from 1740 to 3626. In total, 24,120 genes in 21,750 cells passed the Seurat quality control filtering (see **Methods**) and were used for downstream analysis (**Supplementary Table 1**). The whole transcriptome of all single cells was used to perform differential gene expression and clustering analysis. The result was analyzed by both Principal Component Analysis (PCA) (**Supplementary Figure 5 a-c**) and t-distributed stochastic neighbor embedding (tSNE) (Figure 5a, **Supplementary Figure 5 c-d**). Unsupervised tSNE clusting based on top 1000 highly variable genes suggested that inter-patient heterogeneity was stronger than the global transcriptional state of between tumor cells within the same sample, which was consistent with a previous report(53).

**Figure 5.**
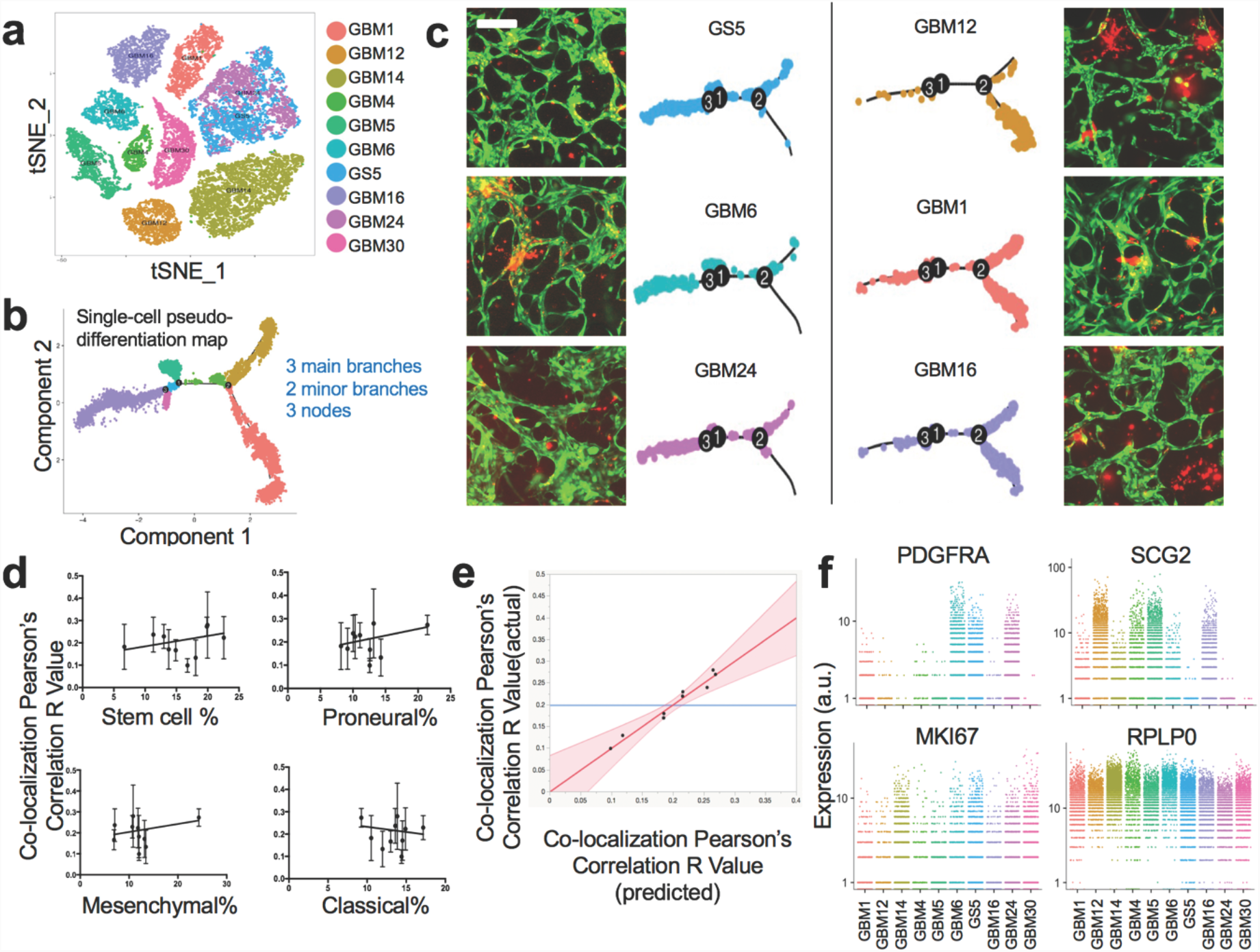
Single-cell RNA-seq correlates on-chip co-localization to transcriptional signatures/subtypes. (**a**) tSNE plot of single cell RNA-seq data from all patient samples. (**b**) Single-cell pseudo-time lineage trajectory obtained by semi-supervised clustering of subtype-specific gene panel using Monocle. (**c**) Representative images of tumor cells in microvasculature and the matched Monocle plots for top 3 and bottom 3 co-localization coefficient samples (Day 4-7). Scale bar: 200 μm. (**d**) The linear regression model showing the percentage of glioblastoma subtype in each sample in correlation with the co-localization R value. (**e**) The multivariate linear mixed model showing the predictor variables (average gene expression of PDGFRA, C1GALT1, THY1, and MKI67) as a combined transcriptional signature that correlates with co-localization. (**f**) Relative gene expression of angiogenesis (PDGFRA, SCG2), proliferation (MKI67), and housekeeping marker(RPLP0) in single cells from all patient samples.

To identify the transcriptional program intrinsic to glioblastoma cells and compare across all ten patients such that the observed *ex vivo* behaviors in the PVN microchip model can be associated with molecular mechanisms, we performed the following informatics analysis. First, we did single-sample Gene Set Enrichment Analysis (ssGSEA)(54) with the gene signatures previously reported(55) to quantify tumor cells expressing stem cell characteristics and found that most patient samples had a high percentage (12.05%-17.49%) of stem-like cells, except for GBM24 (8.77%) and GBM30 (5.79%) (**Supplementary Table 2**). Second, we validated single tumor cell subtypes compared to a permutated data set (permutation=1000) in ssGSEA. According to The Cancer Genome Atlas (TCGA) data(50) published for human glioblastomas, there are four major genotypes: classical, mesenchymal, proneural, and neuronal, each of which has specific transcriptional signatures (684 genes in total). However, at the single-cell level, a patient’s tumor could consist of multiple subtypes, with the leading subtype presumably defining the genotype of bulk tumor(55). At the transcriptional level, a few tumor cells may be enriched for gene markers of two subtypes. Therefore, we defined the subtype of a tumor cell as the dominant gene set with p-value<0.2 (**Supplementary Table 3**). Recent studies(54, 55) suggested the removal of the neuronal subtype due to its overlap with the proneural subtype and suboptimum identification using current gene signatures. In order to further visualize the lineage relationship between tumor cells in each sample, we applied a pseudo-time differentiation trajectory method to analyze all single cells using the gene sets associated with three major subtypes - classical, mesenchymal, and proneural. We then reconstructed the pseudo-time lineage relationships with three subtype markers (555 genes) using semi-supervised analysis in Monocle(56, 57) package. The resulting plot had three major branches and three major nodes connecting them with two minor branches (Figure 5b). The differential gene expression between branches was shown in **Supplementary Figure 6**. Next, we investigated whether tumor/microvessel co-localization (Figure 4d) correlated with tumor cell heterogeneity and subtype. We found that the top three highly co-localized tumor samples (GS5, GBM6, and GBM24) shared a similar cellular trajectory, in sharp contrast with the graph of the three least co-localized tumor samples (GBM12, GBM1, and GBM16) (Figure 5c), indicating a strong correlation between tumor genomic subtypes and *ex vivo* tumor cell dynamics in the microvasculature-on-a-chip system.

Linear regression analysis was performed to compare the percentage of stem-like, classical, mesenchymal, or proneural GBM cells to the co-localization coefficient (Figure 5d). Although none of them reached statistical significance, we observed a slightly positive co-localization with the percentages of stem-like, mesenchymal, and proneurnal cells, but not classical cells. This finding is in agreement with previous reports(43, 58), which studied the role of the PVN in both stem-like cell fate maintenance and vascular track invasion and found mesenchymal cells to be the most invasive subtype. The BTSC PVN model was primarily validated with the proneural subtype(59, 60), but not the classical subtype, which has high probability of EGFR amplification and features enhanced cell proliferation and tumor growth. However, the entire panel of GBM subtype genes (684 genes) did not result in a statistically significant co-localization correlation. Furthermore, we examined specific gene markers with these panels but that were associated more with pro-angiogenesis and the interaction with endothelium (**Supplementary Figure 7**). Based on a multivariate linear mixed model with patient level effect adjusted, we tested if the average gene expression level of marker genes related to the co-localization coefficient. We found 4 genes (PDGFRA, C1GALT1, THY1, and MKI67) were significantly associated with the co-localization coefficient (Figure 5e, **Supplementary Table 4**). Platelet-derived growth factor receptor alpha (PDGFRA), one of the most distinct signature genes for proneural GBM, was found to be expressed highly in the top three ranked tumor cells. This finding further supports our hypothesis that the “homing” of BTSCs in the PVN is well demonstrated in proneural models but not others(59, 60). Endothelial cells in the vessels usually recruit pericytes or mescenchymal cells via PDGF signaling(61) towards vessel maturation. We propose that tumor cells with a high expression level of PDGFRA respond better to PDGF secreted by endothelial cells, and thus show a high vessel co-localization ratio. SCG2, a gene associated with secretory function, was found to negatively correlate with co-localization, except GBM5, which was confirmed as an active extracellular vesicle producer (Figure 4e). GBM30 was found to express low levels of both PDGFRA and SCG2, but showed the highest proliferation and tumor growth capability in mouse xenografts. VEGFA, one of the most important proangiogenic factors, did not show significant differences in most patient samples and no correlation with co-localization coefficients.

These results suggest that ex vivo behaviors of BTSCs in a 3D microvasculature model can recapitulate pathophysiological characteristics, as shown by our model of the PVN. Additionally, the ability to “home” to the PVN of single tumor cells can be associated with transcriptional subtype and correlates with inter-patient heterogeneity. This is the first demonstration that a tissue-engineered 3D microvasculature system can provide a functional niche to assay the dynamics of primary tumor cells derived from patients, opening a new direction for organ-on-a-chip applications.

## Materials and Methods

### Cell Culture

Primary culture of HUVECs were purchased from Yale Vascular Biology and Therapeutics Core. Green fluorescent protein transfected HUVECs (GFP-HUVECs) were commercially obtained (Angio-Proteomie) and cultured in endothelial growth medium EGM-2 (Lonza) with full supplements. HUVECs and GFP-HUVECs between passage 2 and 6 were used in all experiments. No significant difference between HUVECs and GFP-HUVECs in vessel formation ability was observed. Red fluorescent human glioblastoma cells (RFP-U87) and patient-derived glioma stem-like cells (GS5) were kindly provided by Prof. Jiangbing Zhou’s lab at Yale University. RFP-U87 were cultured in DMEM supplemented with 10% FBS. Fresh patient-derived GBM cells were isolated and cultured from GBM surgical specimens. Extensively rinsed tumor specimens were finely minced and placed in DMEM/F-12 medium (Gibco) with 25 unit/ml Papain (Worthington Biochemical Corp). A series of mechanical dissociations was used to obtain a single-cell suspension. Re-suspended cells were cultured in neural basal medium supplemented with B27 (Gibco), FGF (20ng/ml, Peprotech), and EGF (20ng/ml, Peprotech). Brain tumor-derived neurospheres were evident as early as one week after plating.

### Chip Loading and Maintenance

Microvessels formed by HUVECs were co-cultured with glioma cells in a stack of fibrin gel in the 3D cell culture chips (AIM Biotech). The chip, adapted from Prof. Roger Kamm’s design(62), consisted of three parallel channels: one central cell-containing gel loading channel and two lateral medium flow channels. Endothelial cells were seeded in the fibrin gel by introducing HUVECs (2×10^6^ cells/ml) and GS5/U87 (0.5×10^6^ cells/ml) in the 2.5 mg/ml fibrinogen (Sigma) dissolved in serum-free neural basal medium. Thrombin (5 U per 10 mg fibrinogen, Sigma) was then added to convert the soluble fibrinogen into insoluble fibrin strands. Immediately after gentle mixing, the gel (∼10ul) was pipetted into the gel-loading channel of 3D cell culture chips. The samples were placed in a humidified 5% CO_2_ 37°C incubator for 40 minutes, to allow the fibrin to polymerize. 50% EGM-2 media and 50% neural basal medium supplemented with B27, VEGF (50ng/ml), EGF (20ng/ml), and FGF (20 ng/ml) was then added to the two lateral flow channels and was changed every 12 hours.

### Immunofluorescent Staining and Imaging

For live cell tacking, brain tumor stem-like cells GS5 were incubated with Dil cell membrane dye (1:200, Invitrogen) for 40min in a humidified 37°C incubator. For immunofluorescent staining, devices were fixed by 4% paraformaldehyde (ChemCruz) for 20min at room temperature. Primary antibodies were used at 1:100 overnight at 4°C and secondary antibodies were used at 1:1000 for 1 hour at room temperature (**Supplementary Table 6**). The 3D microvessels were imaged using a confocal microscope (Leica DMi8) and deconvoluted by Hyugens Professional software (Scientific Volume Imaging). Unless otherwise stated, all other images were taken with Nikon Eclipse Ti-S microscope and processed with NIS-Elements software (Version 4.20, Nikon Instruments).

### Single Cell 3’ mRNA Sequencing

Our approach for high-throughput single cell mRNA sequencing was based on a closed microwell array chip developed in our lab (Dura *et al*., In Revision). Microwell arrays were used as the platform for co-isolating single cells and uniquely barcoded mRNA capture beads for single cell mRNA capture following lysis. The dimensions of microwells are dictated by the size of mRNA capture beads (∼35 μm); and chosen as ∼45-55 μm in diameter and ∼50 μm in height to ensure accommodation of only a single bead as well as most mammalian cell types. This choice of dimensions also facilitates straightforward removal of beads after mRNA capture either by purging (for closed-environment cell seeding) or centrifugation (for open-surface cell seeding) after inverting the devices. The throughput of the microwell arrays is up to 70,000 wells to be able to sequence ∼1,000-5,000 cells in a single run where the arrays are loaded with a well occupancy rate of 5-10% to minimize dual occupancy (cell duplets). Master wafers for microwell arrays were fabricated using SU-8 negative resist. A single layer of resist (SU-8 2035, MicroChem) was spun at 2200-2400 rpm for 30s to yield feature heights of ∼50 μm. The wafers were then exposed to ultraviolet light through a transparency mask (CAD/Art Services) to pattern microwells. After developing and baking, wafers were hard baked at 150°C for 30 min, and silanized for 2 h in a vacuum chamber saturated with Trichloromethylsilane (Sigma-Aldrich). Fabrication of microfluidic channels followed a similar fabrication procedure using SU-8 2075 where channel height was set to ∼120 μm (1700-1800 rpm for 30 s).

Devices were made by casting polydimethylsiloxane (PDMS, Sylgard 184, Dow Corning) over the master wafers followed by degassing and curing at 80°C for 6-8 hours. Both microwell array and microfluidic channel device were set to a final height of 3-4 mm. After curing, PDMS was peeled off, and devices were cut to proper sizes to fit on a glass slide. For microfluidic channels, holes for fluidic connections were punctured using a biopsy punch (Miltex, 1.5mm). Microwell arrays were first plasma-bonded to microfluidic channel and then to a glass slide. For single cell RNA-seq experiments, the patient-derived GBM neurospheres were cultured for 3∼5 weeks in serum-free media as described above and then gently pipetted with a 200ul pipette tip to dissociate into single cells. Dissociated single cells and mRNA capture beads were then inputted into the microwell chip sequentially using a pipette and allowed to settle into wells by gravity using syringe-pump driven flow. Freeze-thaw lysis buffer was introduced into the microchannel followed by fluorinated oil (Fluorinert FC-40) to seal each microwell to prevent cross-contamination. Cell lysis was achieved using three freeze-thaw cycles, and cell lysate and beads were incubated at room temperature for 60 min for mRNA capture on beads. Following incubation, beads were removed from the microfluidic device by flushing the beads out into an Eppendorf tube. Following bead removal, reverse transcription and library construction were performed as previously described (Dura *et al*., In Revision).

### Fixation and Dehydration of Brain Tumor Stem Cells Near Microvessels for Scanning Electron Microscopy

For the field emission scanning electron microscopy (FE-SEM) observation, we directly fixed and dehydrated the cell-containing gel 7-10 days post-embedding. The transparency film at the bottom of 3D cell culture chip was physically removed by a tweezer. Fixation and dehydration for SEM preparation was conducted using the same process described in our previous paper(63). In brief, the cells were first fixed with 2.5% glutaraldehyde for 1 hour at 4°C, followed by secondary fixing with 1% osmium tetroxide for 1 hour at 4°C. Then, fixed cells were dehydrated by using a grade series of ethanol concentrations (25%, 50%, 75%, and 95%) and followed by final dehydration with 100% ethanol twice for 10 min at 4°C. Subsequently, dehydrated cells were frozen for 3 hours at −80 °C, and then air dried for 24 hours using vacuum desiccator. To observe the cells using FE-SEM, the dehydrated cells were sputter-coated with a layer of iridium (∼10 nm) for obtaining conductive surface on the surface. A FE-SEM (SU-70, Hitachi) was used for observation.

## Data Analysis

### Tumor Morphology and Localization Analysis

The morphology of BTSCs were analyzed with NIS-Elements software and ImageJ. From day 4 to day 6, representative images of each patient BTSCs with GFP-HUVECs were selected for analysis. Each raw image of membrane-labeled fluorescent BTSC cells (Data type: TIFF, size: 1392×1940 pixels^2^, about 0.12 mm^2^) was binarized by the auto threshold tool and manually segmented through freehand tool. Co-localization correlation of tumor cells and endothelial cells was evaluated by the Pearson’s correlation coefficient(64) by the co-localization test tool in Fiji software(65). The coefficient ranges from −1 to 1: +1 for perfect correlation, 0 for no correlation, and −1 for perfect anti-correlation.

### Cell Motility

GFP-HUVECs and Dil-stained patient-derived GS5 cells in fibrin gel were seeded in the AIM chip and allowed to grow for 48 h. After pre-mature vessel formation, the plate holder of the chip was mounted on a Nikon Eclipse Ti-S microscope with a motorized stage (Prior Scientific) and an environment control incubation chamber (Okolab) to maintain 37°C with 5% CO_2_. Phase contrast and fluorescent images were recorded every 30 min for 20 h using a CCD camera (ANDOR) with a low magnification 4X Fluor objective. Each single cell was manually labeled in the continuous frames for 20 h. Cell motility parameters were assessed via tracking of single tumor cells (n=3, ∼20 cells per cropped region per trial) using Fiji. Motility was defined as the distance traveled in a unit time. Cellular displacement was calculated using the corresponding x and y coordinates at initial time t_0_ and end time t_n_. Trajectory was plotted by Matlab (R2017a, MathWorks).

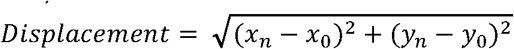

### Finite Element Simulation (FEM) Analysis

COMSOL Multiphysics® (Version 5.0, COMSOL) software was used to perform finite element analysis on the AIM Biotech microfluidic device set up. 2D fluorescent images of microvessels formed in the AIM Biotech devices were created in COMSOL and simulations were performed to mimic the microenvironment the vessels and cells were experiencing. A whole chip scan image of GBM6 in the microenvironment of green fluorescent HUVEC microvessels (Day 4) was first bianarized using ImageJ. Then, using WinTopo Raster to Vector Conversion Software (v1.76, SoftSoft Ltd.), the black and white BMP images were converted to vectorized images which could be imported into COMSOL. To successfully vectorize the images in WinTopo, various image processing techniques such as erosion, despeckle, and prune were used to create an acceptable image that could be processed using edge detection. The vector file was then imported into COMSOL and used to reconstruct the microvessel environment. The free and porous flow module was applied to simulate for media and fibrin respectively. Media flow with a density of 1020 kg/m^3^ and viscosity of 0.8cP was governed by the Navier-Stokes and continuity equations for laminar flow(66, 67). The fibrin was modeled as a porous matrix with a porosity of 0.99 and a permeability of 1.5×10^−13^ m^2^ and was simulated using the Brinkman equation(68). The geometry was scaled and meshed in COMSOL with a minimum element size of 0.09mm and a maximum element size of 0.6mm. For initial conditions, a pressure gradient of 10Pa from the top to bottom channels was used to represent a column height difference of 1mm. Hydrostatic pressure from a column height of 1mm representing 10Pa was used as the inlet pressure while the outlet pressure was set to 0 Pa.

### Single Cell mRNA Sequencing Analysis

In total, we sequenced 26,027 single cells by using 75 bp pair-end reads on a HiSeq2500 instrument (Illumina) in HighOutput Mode V4. Raw reads were preprocessed for cell barcodes and UMIs, and then aligned to the human genome(hg19) using STAR v2.5.2b as described in Dropseq method(69). Digital expression matrix was generated for the cells with over 10,000 reads per cell.

The Seurat(70) package (V2.3.0) in R (V3.4.1) was applied to identify differentially expressed genes among 26,027 single cells from nine different GBM patients and one GBM cell reference (GS5). Cells were considered in the analysis only if they met the following quality control criteria: 1) expression of more than 1,000 genes and fewer than 5,000 genes; 2) low expression of mitochondrial genes (<10% of total counts in a cell). After filtering, 24,120 genes in 21,750 cells were left for clustering analysis. Genes that were differentially expressed in each cluster were identified using the Seurat function FindMarkers, which returned the gene names, average log fold-change, and adjusted p-value for genes enriched in each cluster. Unsupervised clustering in principal component analysis (PCA) was performed with 30 statistically significant principal components that were identified from the top 1000 highly variable genes from all the samples. We then projected single cells onto a two-dimensional map using t-Distributed Stochastic Neighbor Embedding (t-SNE) to discover inter-patient heterogeneity.

The Monocle(56, 57) package (V2.6.4) was used to plot single cell pseudo-time trajectories to discover the behavioral similarity and transitions. We use the proneural, mesenchymal, and classical subtype genes identified before(50) to perform the semi-supervised analysis. Monocle looked for variable genes and augmented the markers when construct the clustering and ordering of the cells. DDRTree algorithm was used to visualize the pseudo-time trajectory in the reduced dimensional space. Plot_genes_branched_heatmap module was applied to plot out the genes (qval<1e-300) that had similar expression profile on a branch.

## Statistics

Results are shown as mean +/- standard deviation. Student t-test is used to assess the comparisons between the groups in Figure 2. Statistical analysis (One Way ANOVA) demonstrated significant difference among three groups of these measurements in Figure 3d-e. Statistical significance was assumed for p<0.05, unless otherwise specified. In figure 5e, a multivariate linear mixed model was performed with patient level effect adjusted. Akaike information criterion (AIC) and Bayesian information criterion (BIC) were utilized for model selection. All tests were performed with Prism (Version 7.0, GraphPad Software), JMP (Version 13.0, SAS Institute), or R (V3.4.1).

## Sequencing Data Availability

The single-cell RNA-seq data will be deposited in the Gene Expression Omnibus(GEO) following the acceptance of the manuscript.

## Supplemental Materials

Supplemental information includes 7 figures, 9 tables, 3 videos, and supplemental text.

## Author Contributions

Y.X. and R.F. conceived the project and designed the experiments; Y.X., D.K., B.D., K.Z. performed the experiments; J. L., P.Z., and J. Z. prepared cells from the patient tissue; Y.X., R.Y., H.L., E.H., J.I., and A.C. analyzed the data; R.F. and J.Z. supervised the project; Y.X. and R.F. wrote the paper.

## Acknowledgments

We thank Drs. Laura Niklason, Eric Holland, Franziska Michor, and Frank Szulzewsky for scientific discussion. We thank Misha Guy, Vladimir Polejaev, Zhenting Jiang, and Alice Yun for suggestions and help on the simulation computing and SEM/confocal imaging process. This research was supported by the Packard Fellowship for Science and Engineering (R.F.), National Science Foundation CAREER Award CBET-1351443 (R.F.), U54 CA193461 (R.F.), U54 CA209992 (Sub-Project ID: 7297 to R.F.), R01 NS095817 (J.Z.), Yale Cancer Center Co-Pilot Grant (to R.F.). The molds for microfluidic devices were fabricated in the Yale School of Engineering and Applied Science cleanroom. Sequencing was performed at the Yale Center for Genome Analysis (YCGA) facility. Data was analyzed at Yale High Performance Computing (HPC) center. Super resolution confocal imaging was performed at Yale Center for Cellular and Molecular Imaging (CCMI).

